# Naturally cysteine-less LOV domains from halophilic archaea exhibit magnetic field effects on their fluorescence

**DOI:** 10.64898/2026.07.30.741865

**Authors:** Brian L. Ross

**Affiliations:** Quantum Biology Institute, Los Angeles, USA

## Abstract

Magnetic fields can modulate the function of certain flavoproteins through the radical pair mechanism, in which they influence the spin evolution of a coherent pair of radicals. This process can result in magneto-fluorescence, in which magnetic fields modulate the intensity of fluorescence emitted from the flavin cofactor. The prevalence of this property across protein families and domains of life, however, remains poorly characterized. In canonical Light-Oxygen-Voltage (LOV) domains, a conserved cysteine forms an adduct with the flavin that leads to downstream signaling. Mutating this cysteine instead yields signaling through the neutral semiquinone radical form, and the same mutation was crucial for enhancing magneto-fluorescence in the engineered protein MagLOV2. We therefore hypothesized that natural LOV domains that lack this conserved cysteine may exhibit magneto-fluorescence. Using a custom magneto-fluorescence imaging platform, we measured the fluorescence of E. coli colonies expressing three such domains, as well as a single mutant of one of them, under a switched external field. HsuLOV, from a halophilic archaeon, exhibited magneto-fluorescence, as did a single mutant of BAT-LOV, from a second halophilic archaeon. The domain amb2291 from magnetotactic bacteria, on the other hand, showed no detectable response under our illumination conditions. Magneto-fluorescence is therefore not exclusive to engineered proteins but is present in natural cysteine-less LOV domain sequences. The proteins reported here are, to our knowledge, the first magnetosensitive proteins reported from archaea. This suggests that radical-pair magnetosensitivity may be more widespread across LOV domains than previously appreciated.

## 1. Introduction

A handful of flavoproteins have previously been shown to exhibit biological functions that are modulated by relatively weak magnetic fields. The leading explanation for how these proteins sense weak magnetic fields is the radical pair mechanism (RPM), in which electron transfer generates a pair of spin-correlated unpaired electrons [1]. The coherent evolution of their spin state can be influenced by an external magnetic field, altering the relative populations of singlet and triplet states and, consequently, the chemical reaction and macroscopic outcome. While evidence supporting the RPM has grown substantially in recent years, the molecular determinants of protein magnetosensitivity, as well as the prevalence of this phenomenon both across the tree of life as well as within and across protein families, remain poorly characterized.

For historical reasons, the RPM has been studied most extensively in cryptochrome, a protein important for circadian rhythm control that is also hypothesized to enable migratory birds to sense and follow the magnetic field of the Earth [2]. Magnetic field sensing has also been described in the DNA repair enzyme photolyase [3], redox enzymes [4], as well as Light-Oxygen-Voltage (LOV) domains [5–7]. Recently, researchers engineered a fluorescent protein with an enhanced magnetic response called MagLOV (and its variants MagLOV2 and MagLOVf), derived from a plant LOV domain, through multiple rounds of directed evolution [5, 6]. The resulting protein exhibits a large magnetic field effect (MFE), meaning that its fluorescence intensity changes appreciably when exposed to different magnetic field conditions.

Canonical LOV domains are activated by blue light, in which photoexcitation of their flavin cofactor initiates the formation of a covalent bond with a nearby conserved cysteine residue, forming an adduct. This event induces a conformational change in an adjacent conserved glutamine residue, reorganizing the local hydrogen-bonding network and propagating structural changes that enable downstream signaling [8]. However, Yee *et al*. demonstrated that signaling can still occur even when the conserved cysteine is mutated [9]. Instead of forming the canonical cysteinyl adduct, these cysteine-less mutants photoreduce the flavin to the neutral semiquinone (NSQ) radical state. Signaling activity is nevertheless retained provided the conserved glutamine remains intact, indicating that formation of the adduct is not required for signal transduction. Formation of the NSQ state proceeds through a spin-correlated radical-pair intermediate, in conjunction with an electron donor, that can, in principle, exhibit magnetic-field sensitivity via the RPM. Indeed, mutation of the conserved cysteine was central to the development of MagLOV and its variants, as it redirects photochemistry toward the NSQ-forming pathway capable of being modulated by magnetic fields.

The MFE of MagLOV2 emerges at relatively low magnetic fields and saturates at approximately 60 mT, with spin-state mixing driven by hyperfine asymmetry between a radical pair formed between the FMN and a tryptophan residue. On the other hand, it was recently demonstrated that AsLOV2, the wild-type LOV photoreceptor from which MagLOV was engineered and which retains the conserved adduct-forming cysteine, exhibits a high-field MFE that emerges above 100 mT [7]. Unlike MagLOV2, this effect is thought to arise from a radical pair between FMN and the conserved cysteine, in which an unusually large difference in the radicals’ g-factors drives spin-state mixing at high field strengths. While this recent work established that natural LOV domains can exhibit high-field MFEs, it remains unknown whether naturally occurring LOV domains also display the low-field onset MFEs, similar to MagLOV and its variants.

Recently, three naturally cysteine-less LOV domains from bacteria and archaea have been described and optically characterized, outside of the context of magneto-fluorescence. Yee *et al*. demonstrated that a naturally cysteine-less LOV domain from the bacterio-opsin activator protein taken from the halophilic archaeon *Halorubrum hochstenium* (BAT-LOV) was capable of signal transduction [9]. Furthermore, a naturally cysteine-less LOV domain called amb2291 from the magnetotactic bacterium *Magnetospirillum magneticum* has been shown to be involved in oxidative stress adaptation, regulation of magnetosome-related gene expression, and the coordination of phototactic and light-dependent magnetotactic behaviors [10, 11]. Lastly, a LOV domain from the halophilic archaeon *Halanaeroarchaeum sulfurireducens* (HsuLOV) was demonstrated to reversibly photoreduce to the NSQ, especially in high salinity and low oxygen environments [12].

Because these naturally cysteine-less LOV domains signal through the NSQ radical form, they have the potential to participate in the RPM and magnetosensing, similar to MagLOV and its variants. In this study, we characterize these three naturally cysteine-less LOV domains’ fluorescent responses to external magnetic fields.

## 2. Methods

### 2.1. Protein expression in E. coli

Sequences for the LOV domain core plus J*α* helix were chosen for HsuLOV (accession AKH96799.1, residues 266-399), amb2291 (accession WP_231848828.1, residues 335-467), and BAT-LOV (accession WP_008581736.1, residues 141-274). E. coli codon-optimized DNA constructs for these protein sequences (as well as a W172F variant of BAT-LOV), cloned into the bacterial expression plasmid pET28a, were ordered from Twist Biosciences. These constructs were transformed via heat shock into BL21(DE3) *E. coli* and plated onto LB agar plates supplemented with 50 μg/mL kanamycin. These plates were incubated at 37°C overnight, after which they were transferred to room temperature for 24 hours in the dark before imaging.

### 2.2. Magneto-fluorescence Bacterial Plate Imaging

A custom-built magneto-fluorescence imaging platform was used to image bacterial plates under controlled magnetic field sequences, as previously described [13, 14]. Fluorescence images were acquired every 2 s using 470 nm excitation light at an illumination intensity of 6.6 mW/cm^2^. During each imaging cycle, the sample was illuminated for 1.6 s, followed by a 400 ms camera exposure. An electromagnet switched the magnetic field on and off every 90 s, with the ON state corresponding to a 100 mT magnetic field applied in the downward direction. Additional details of the instrument and its construction are available in our Electronic Lab Notebook [15].

### 2.3. Data analysis

Fluorescence images of bacterial plates were processed using the custom-developed mfe-fit software package, which is freely available in a GitHub repository [16]:, as previously described [14]. Images were automatically segmented into colonies using rolling ball background subtraction, followed by thresholding with the Otsu algorithm with watershed separation. For each colony, the mean fluorescence intensity was measured and the background fluorescence from a nearby region was subtracted.

To isolate the MFE, we estimated the fluorescence trace expected in the absence of an applied field using PCHIP (Piecewise Cubic Hermite Interpolating Polynomial) interpolation through anchor points drawn from the field-OFF segments [17]. Nine anchors were placed across the first OFF segment, which had no ON segment preceding it, so the entire segment was considered a valid baseline. Because fluorescence requires time to recover to baseline once the field is switched off, each subsequent OFF segment contributed a single anchor, obtained by fitting a line to its last five frames and evaluating it at the acquisition time of the final frame. The normalized residual fluorescence (*r*(*t*)) was estimated via the expression:

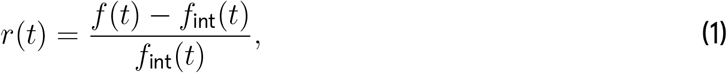

where *f* (*t*) is the fluorescence trace and *fint*(*t*) is the interpolated fluorescence baseline. This represents the deviation from the baseline due to the repeated application and removal of the external magnetic field. The MFE for each ON segment was computed as the normalized residual of the final five frames, corrected by subtracting the normalized residual of the final five frames from the preceding OFF segment. To aggregate the data, the MFEs from OFF-ON cycles were averaged within each colony trace to calculate a per-colony mean. We excluded the first OFF–ON cycle, where an initial transient drop in fluorescence may bias the baseline estimate. Finally, the per-colony means were averaged across colonies within a plate.

## 3. Results

To investigate the effect of external magnetic fields on the fluorescence intensity of naturally cysteine-less LOV domains, E. coli colonies expressing HsuLOV, amb2291 and BAT-LOV were imaged using our custom magneto-fluorescence imaging platform. A W172F mutation in BAT-LOV had previously been shown to increase fluorescence lifetime, intensity and photoreduction rates [9]. Therefore, we also tested BAT-LOV W172F for magneto-fluorescence in addition to wild type (WT).

HsuLOV demonstrated magnetosensitivity in its fluorescence intensity, with a decrease in fluorescence observed upon applying a 100 mT field that recovered upon the removal of said field (Fig. 1a). amb2291, on the other hand, did not show such a clear response, which, if present, was below the noise (Fig. 1b). However, MFEs in fluorescence are highly dependent on the illumination intensity, and thus the lack of observed MFE in our particular experimental setup does not rule out magnetosensing in amb2291 (see Discussion).

**Fig. 1.**
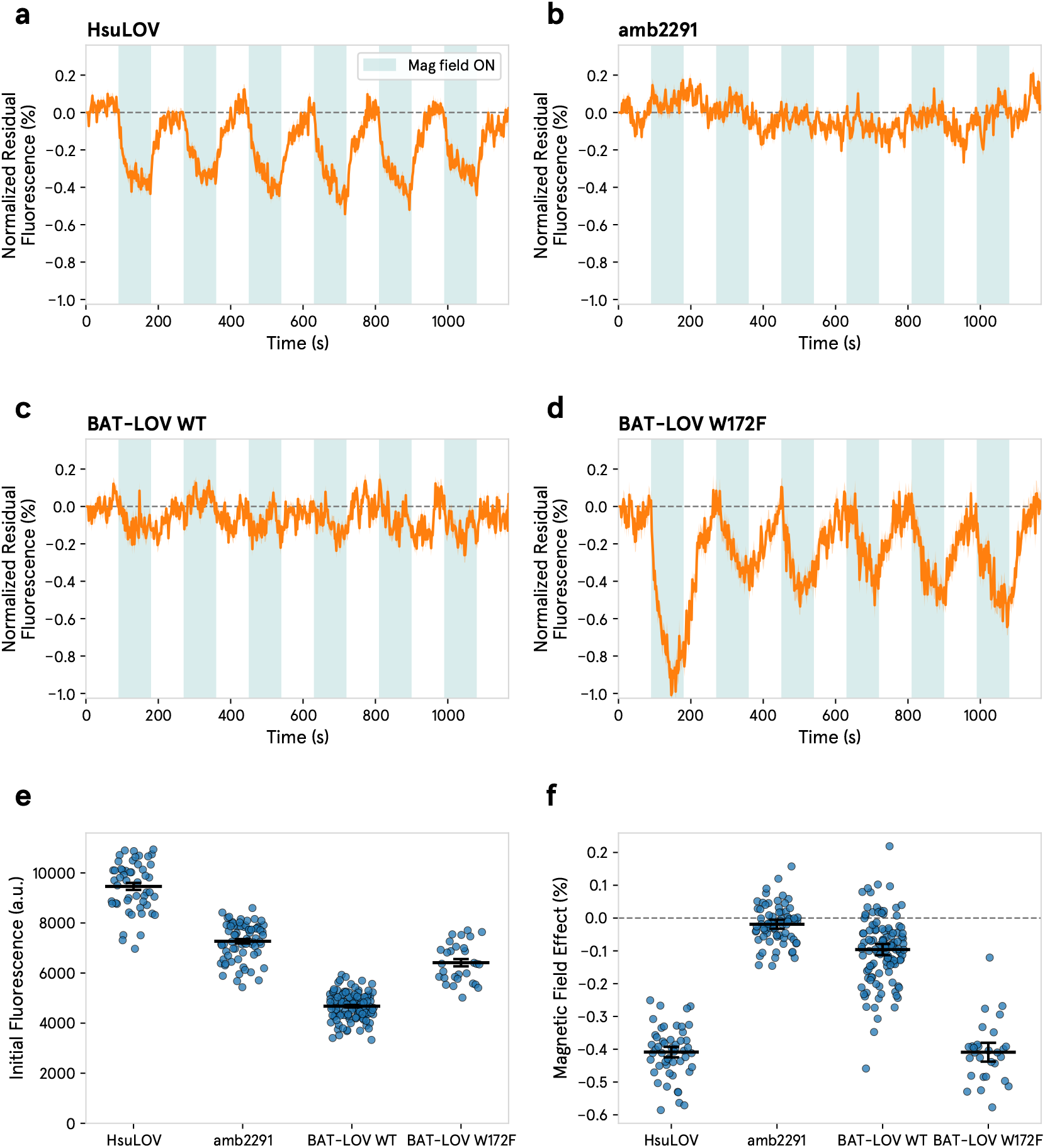
Magnetosensitivity is observed in naturally cysteine-less LOV domains from archaea. Mean traces of normalized residual fluorescence for E. coli expressing a) HsuLOV (n=51), b) amb2291 (n=71), c) BAT-LOV wild type (WT) (n=118), and d) BAT-LOV W172F (n=29). Teal shading indicates periods with the 100 mT field applied. e) Initial fluorescence intensity for each construct. f) Per-colony MFEs, taken as the mean across external magnetic field OFF-ON cycles, excluding the first cycle. In e) and f), each dot is one colony; the black bar is the mean. In e), error bars are the standard error of the mean across colonies, while in f), error bars are the combined uncertainty (see Methods).

The fluorescence of BAT-LOV WT, likewise, was at most weakly magnetosensitive in the present setup (Fig. 1c), and an MFE was not discernible by visual inspection of the fluorescence trace.

However, the BAT-LOV W172F mutant showed magnetosensitivity of its fluorescence intensity in our setup (Fig. 1d), with a clear decrease in fluorescence upon application of the external magnetic field, and corresponding recovery when the field was removed. A larger response was observed in the first cycle, but that may be due to inaccuracies in the baseline estimate during an initial fluorescence transient at the beginning of the experiment. The mutation, consistent with what had been previously reported, did result in a higher measured initial fluorescence intensity as compared with BAT-LOV WT (Fig. 1e) [9].

Fig. 1f shows the MFEs measured for each colony. The magnetosensitivity of the fluorescence of HsuLOV and BAT-LOV W172F, as captured by the present setup, is unambiguous, with average MFEs of -0.409% ± 0.016% and -0.409% ± 0.029%, respectively. BAT-LOV WT showed an average MFE of -0.096% ± 0.017%, with per-colony values skewed towards negative deviations, suggestive of a weak effect, even if it is not obvious from the mean fluorescence trace. amb2291 did not show any measurable MFE in the present setup.

## 4. Discussion

Together these data demonstrate that magneto-fluorescence is a property found in natural protein sequences and does not require engineering or directed evolution. Furthermore, to our knowledge, no magnetosensing proteins had yet been identified in archaea, so the demonstration of magnetosensitivity of HsuLOV reflects the first demonstration of magnetic field effects in proteins from the archaeal domain. Together with magnetosensitive proteins previously described in bacteria and eukaryotes, these results indicate that protein magnetosensing is a phenomenon found across all domains of life.

Based on a theoretical understanding of the RPM and knowledge of MagLOV2’s response to magnetic fields, we were able to hypothesize that naturally occurring LOV domains that lack the conserved adduct-forming cysteine residue might be magnetosensitive, which we confirmed through magneto-fluorescent imaging. Unlike MagLOV2, however, these cysteine-less LOV domains do maintain the conserved glutamine, which is important for LOV domain activity. Therefore, we further predict that these naturally occurring LOV domains may be able to induce conformational changes and transmit signaling in a magnetically dependent fashion, which would be useful for developing genetically encodable, magnetically actuated molecular tools. Furthermore, while the magnitude of the MFE of HsuLOV and BAT-LOV magneto-fluorescence is modest, based on the success of MagLOV2 engineering, it is likely that the magnitude and kinetics of the response could be modulated and optimized through directed evolution.

Because these proteins lack the adduct-forming cysteine, we hypothesize that their MFEs are low-field onset, similar to MagLOV2, rather than the high-field effect observed in wild-type AsLOV2. However, because the present study examined fluorescence at only 100 mT, we cannot determine the field dependence of the MFE and therefore cannot exclude a high-field onset mechanism. Future characterization of the relationship between fluorescence intensity at different magnetic field strengths will distinguish between these two possibilities.

A number of different biophysical factors can contribute to protein magneto-fluorescence.

The sign of the MFE is determined both by the strength of the field and the starting spin state of the electron pair. Because we used a field strength (100 mT) that is much stronger than the typical hyperfine interactions, the fact that the MFEs for HsuLOV and BAT-LOV W172F were negative suggests that the radical pair is born in the triplet state, meaning that intersystem crossing is faster than electron transfer [18]. Moreover, for a protein to sense a magnetic field via the RPM, the flavin radical must be stable, and the coherence of the superposition of the radical pair must be maintained for long enough for the magnetic field to have an appreciable effect (at least on the order of the period of Larmor precession). In addition, the orientation of potential electron donors and the flavin acceptor must be suitable for electron transfer. Finally, the local magnetic field environments of the two radicals, determined by the nuclear spins that they interact with, must be sufficiently different to induce evolution of spin states. It is possible that the differences in magneto-fluorescence we observe between the cysteine-less LOV domains in this study, and other magnetosensing proteins in general, are due to variations in any of these factors.

However, it is important to note that even if a flavoprotein can sense magnetic fields through the RPM, that does not mean that it will exhibit experimentally measurable magneto-fluorescence. Therefore, just because amb2291, for example, did not exhibit a measurable MFE in its fluorescence trace on our particular experimental setup does not mean that the protein does not function in a magnetic-field dependent way. Our previous work in kinetic modeling of flavoprotein fluorescence demonstrated that whether or not an MFE is experimentally observed through fluorescence intensity readouts depends on the relative rates of two different processes: 1) photoexcitation and 2) the spin-independent chemical reactions, such as protonation/deprotonation and redox processes, that recover the flavin ground state after the loss of coherence of the radical pair [19]. Fluorescence contrast between singlet and triplet states is caused by the fact that singlets can undergo back electron transfer to recover the ground state and feed back into the photocycle, which creates the measured signal, while triplets are spin-forbidden to do so. However, if the illumination intensity is too low, such that photoexcitation is the rate limiting step and not the recovery of the ground state, then it is impossible to experimentally distinguish the fluorescence resulting from the different radical pair spin states. Therefore, differences in magneto-fluorescence between the different LOV domains tested in our setup may not actually be a result of differences in magnetosensitivity itself, but may also be due to different chemical properties of the flavin binding pocket, which affect the rates of the redox and protonation/deprotonation reactions that recover the ground state flavin and thus alter the kinetic rates of the photocycle. Consequently, the absence of a measurable fluorescence MFE under our experimental conditions does not necessarily indicate the absence of radical-pair-mediated magnetosensitivity. The illumination intensity used here, 6.6 mW/cm^2^, is relatively low, so only proteins for which the reactions which recover the ground state are relatively slow would exhibit measurable magneto-fluorescence in our experimental setup.

## 5. Conclusion and Future Directions

This study is the first demonstration of magnetically modulated fluorescence in naturally occurring LOV domains that lack the conserved cysteine as well as of magnetosensing proteins in the archaeal domain of life. We were able to leverage our understanding of the RPM and previous experience with engineered MagLOV proteins to predict that naturally occurring cysteine-less LOV domains may exhibit magnetosensitive fluorescence.

Future work will include characterizing the relationship between fluorescence and magnetic field strength in these naturally cysteine-less LOV domains to conclusively determine whether they exhibit the low-field onset MFEs characteristic of MagLOV2 or the high-field onset effects recently reported for AsLOV2. We will also perform phylogenetic analyses of bacterial and archaeal LOV domains to identify additional cysteine-less proteins. Furthermore, we aim to engineer these noncanonical cysteine-less LOV domains for expanded dynamic range and altered kinetics in order to create alternative magneto-fluorescent proteins that retain functional signaling. Because these domains retain the conserved glutamine residue, we also aim to determine whether they undergo conformational changes and transmit signals in a magnetic field-dependent manner. Additionally, we seek to understand how these LOV domains function in their native contexts and determine whether their magnetosensitivity has been evolutionarily selected for or is simply an incidental consequence of their photochemistry. Ultimately, we hope to leverage these properties to develop magneto-actuated molecular tools.

## 6. Author Contributions

Following the CRediT taxonomy [20]:

1. Brian L. Ross: conceptualization; data curation; formal analysis; investigation; software; visualization; writing – original draft

## 7. Acknowledgements

I would like to thank Morgan Sosa for her assistance with formatting, revising, and editing the manuscript, and I thank Venkatesh Sridharan and Clarice D. Aiello for critically reading the manuscript and providing helpful feedback. I would like to thank Alessandro Lodesani for his help with instrumentation.

